# Uncovering the evolutionary tail of GZMM: An NSP4 related protease

**DOI:** 10.1101/2022.06.02.494303

**Authors:** Ahmer Bin Hafeez, Jamshaid Ahmad

## Abstract

Serine proteases are the most predominant class performing a number of activities in organisms. Undergoing several mutations in their sequence over a span of a billion years yet S1 chymotrypsin/trypsin family has maintained a common fold. Granule Associated Serine Peptidases of Immune Defense (GASPIDS) belonging to the S1 class, found in the granules of immune cells are explicitly involved in the regulation of immune-related functions possessing a conserved catalytic triad Ser-Asp-His. The neutrophils along with other cells express four serine proteases (ELA2, PR3, CTSG and NSP4) sharing certain common characteristics. Similarly, CTLs and NK cells express a set of proteases, Granzymes. This study infers an evolutionary relationship among GASPIDs. We employed computational strategies and found that a higher degree of similarity existed between NSP4 and GZMM as compared to their members i.e. NSPs and granzymes, respectively. Similarly, GZMM a protease of NK cells and t cells lineage is found in the Met-ase locus consisting of NSPs genes i.e., *Ela2, Prtn3* and *Ctsg*. The evolutionary relationship of *Prss57*/NSP4 and *gzmm*/GZMM was reconstructed through empirical phylogenetic analysis which revealed *Prss57*/NSP4 to be closely related to *gzmm*/GZMM. Additional co-expression analysis was carried out to determine the regulatory role of *Prss57*, inducing *Gzmm*. From this work, we inferred that *Prss57*/NSP4 is closely related to *Gzmm*/GZMM.

## Introduction

Serine proteases constitute one of the largest groups of proteases [1,2], given the name due to the presence of a serine amino acid in their active site, which is key in performing catalysis [3,4]. Serine along with histidine and aspartate makes the active site also referred to as the catalytic triad that has remained conserved among this class of proteases across the species [4,5]. Other serine proteases with peculiar catalytic dyad and tetrad have also been identified, analogously comprising serine as a catalytic residue [6].

There are over 26,000 serine proteases [7], classified into 13 clans and 85 families [8]. This classification system was put forward by Barrett and Rawlings [9] who classified the proteases on the basis of their sequence similarities, catalytic properties and structure, integrating them in a database named MEROPS [9–11]. Brenner was the first to report TCN and AGY codons for serine and argued enzymes in the same family coded by different active site codons belong to a different lineage because the interconversion of TCN↔AGY is very rare [12]. The serine proteases of the S1 family (trypsin/chymotrypsin) have undergone sequence changes but still maintained a common fold over an evolutionary period of more than one billion years [13]. However, the diverse functionalities of this family are attributed to the C-terminal region in the protease domain. It is due to the fact that the C-terminal portion in serine proteases constitutes most of the area of S1-S3 specificity sites [13]. Moreover, the majority of the residues that shield the access to the active site are found in the C-terminal tail sequence. The C-terminal constituting residues comprises 95% of the area encompassing the primary substrate specificity site S1. The active site Ser195 and residue 225 that are essentially involved in S1 site architecture along with the residues 216 and 226 that controls access to the active and primary S1 site are also found in this region. Whereas, 70% of the area around the around S2 and S3 substrate specificity is also covered by the C-terminal site [13]. The known prominent roles of serine proteases in the mammalian species are immunological and non-immunological. The serine proteases involved in immune defense i.e. apoptosis, cytokine & inflammatory responses and direct killing of pathogens via phagosomes [2,7,14], are more commonly known as GASPIDs (Granules Associated Serine Peptidases of Immune Defense) [2] Whereas, the non-immunological serine proteases involved in digestion, blood coagulation, fibrinolysis, development, and fertilization are recently been referred to as non-GASPIDS [14].

GASPIDs are synthesized by a number of immune cells. Neutrophils, being the most abundant cells of the immune system, synthesize, in their azurophilic granules, a group of four structurally similar serine proteases collectively known as neutrophil serine proteases (NSPs) [15]. These neutrophils serine proteases are Neutrophil Elastase (NE), Cathepsin G (CTSG), Proteinase 3 (PR3) and Neutrophil Serine Proteases (NSP4) [15]. These proteases are active against a wide variety of bacteria and fungi, inactivating and killing them [16]. Similarly, mast cells are predominantly involved in allergic responses and confer immunity against parasites through the synthesis of mast cell chymase and mast cell tryptase [17,18]. The cytotoxic T lymphocytes synthesize a variety of immune serine proteases known as granzymes that initiate apoptosis in the virally infected and tumorous cells. Granzymes identified so far are granzyme A, B, H, K and M [19,20].

NSP4, the fourth member of NSP is described recently with an Arg-like specificity sharing approximately 40% sequence identity with the three previously identified NSPs [21].

NSP4 is encoded by the gene *Prss57* located on the P arm of chromosome number 19 in humans (19p13.3) and on chromosome 10 in mice inside the Met-ase locus that comprises trypsin-like serine proteases granzyme M, PR3, NE, complement factor D and azurocidin [20,21]. NSP4 is proteolytically processed by dipeptidyl-peptidase I (Cathepsin C) that cleaves a short propeptide sequence at the N-terminus to form a functionally active mature form of NSP4 [15,21]. NSP4 is found to be highly conserved among the vertebrate species from bony fish to humans and originated earlier than the other NSPs. Therefore, NSP4 is considered an ancestral protease and must play important role in neutrophil biology [22]. GZMM, also known as Met-ase or Met-1 was first identified in the NK cells of mice [23] and subsequently in humans [24]. The gene *Gzmm* is also found in the Met-ase cluster [25]. GZMM as compared to the other granzymes has a propeptide sequence of 6 amino acids whereas others have a propeptide of 2 amino acids [24,26]. Several investigations inferring the evolutionary pattern of serine proteases have been conducted previously [27]. Whereas, recently Akula et al conducted a detailed evolutionary and diversification study on serine proteases [28]. However, both proteases NSP4 and GZMM are expressed by cells in different immune lineages. NSP4 is secreted by neutrophils that belong to myeloid lineage [29]. While GZMM is secreted by the NK cells of myeloid lineage and also by t-cells of the lymphoid lineage [30,31]. The presence of GZMM the only granzyme present in the NSPs coding in Met-ase locus piqued our interest. The purpose of this work was to study the evolutionary history of GZMM. Given that NSP4 is considered an ancient protease, we assume certain *Prss57* gene duplication events may explain the origin of *Gzmm*.

## Methodology

### Sequence identification and retrieval

Ensembl [32] and NCBI [33] databases were probed for the human *Prss57, Ela2, Ctsg, Prtn3, GzmA, GzmB, GzmH, GzmK, and GzmM* genes. The protein sequences of the selected genes NSP4, HNE, CTSG, PR3, GZMA, GZMB, GZMH, GZMK, and GZMM were also downloaded for further study.

The protein downloaded sequences were analyzed by aligning only the mature protein sequences as they are synthesized as pre-pro proteins and require proteolytic preprocessing. Bioedit software was used for aligning and the sequences were aligned using ClustalW algorithm with default parameters [34,35]. The active site and substrate specificity conferring residues were analyzed [36] to look for variation and conservation. The percentage similarity among the protein sequences was calculated using the blastp tool, keeping the parameters default [37].

### Sequence Logo

To find out the conserved regions in both the proteases GZMM and NSP4, a sequence logo was generated, using the default parameters. It is a tool used to identify conserved patterns in a nucleotide or protein sequence. The multiple sequence alignment file of GZMM and PRSS57 in Fasta format was submitted to the WebLogo server available online (http://weblogo.berkeley.edu/logo.cgi).

### Locus Map generations

The genomes were accessed using Ensembl and NCBI. The locus maps for *Prss57* and *Gzmm* were generated and the flanking genes upstream and downstream were observed. Specie specific searches were performed to cull the genes information.

### Phylogenetic analysis

A Phylogenetic tree for the mature protein sequence was constructed using a distance matrix file (supplementary file 1) generated from the structome database (http://structome.bii.a-star.edu.sg/ver2/). The distance matrix file was then imported in splits tree software and neighbor-joining algorithm was used for tree construction. Whereas the C-terminal tail of proteases and codons sequences were aligned using default ClustalW algorithm in bioedit with default parameters. The phylogenetic tree was constructed in MEGAX [38]. Neighbor-joining algorithm [39]was used and the number of bootstrap replication was set to 1000.

### Co-expression analysis of *Prss57* and *Gzmm*

The co-expression and protein-protein interaction analysis of *Prss57* and *Gzmm* were performed using GeneMania (available online and desktop version Cytoscape) to observe the common expressing and co-expressing genes [40]. An interaction network for *Prss57* and *Gzmm* was generated with the cut-off values set at default.

## Results

### Sequence analysis and Alignment

The sequence alignment of NSP4 and GZMM other GASPIDs revealed conservation across previously determined [insert reference] residues of the active site. PHRSYMA sequence was also found conserved. However, variations were observed in substrate specificity conferring residues 189, 190, 192, and 226 of both proteases shown in figure 2. While Ser at position 214 in GZMM and NSP4 is conserved.

**Table 1.**
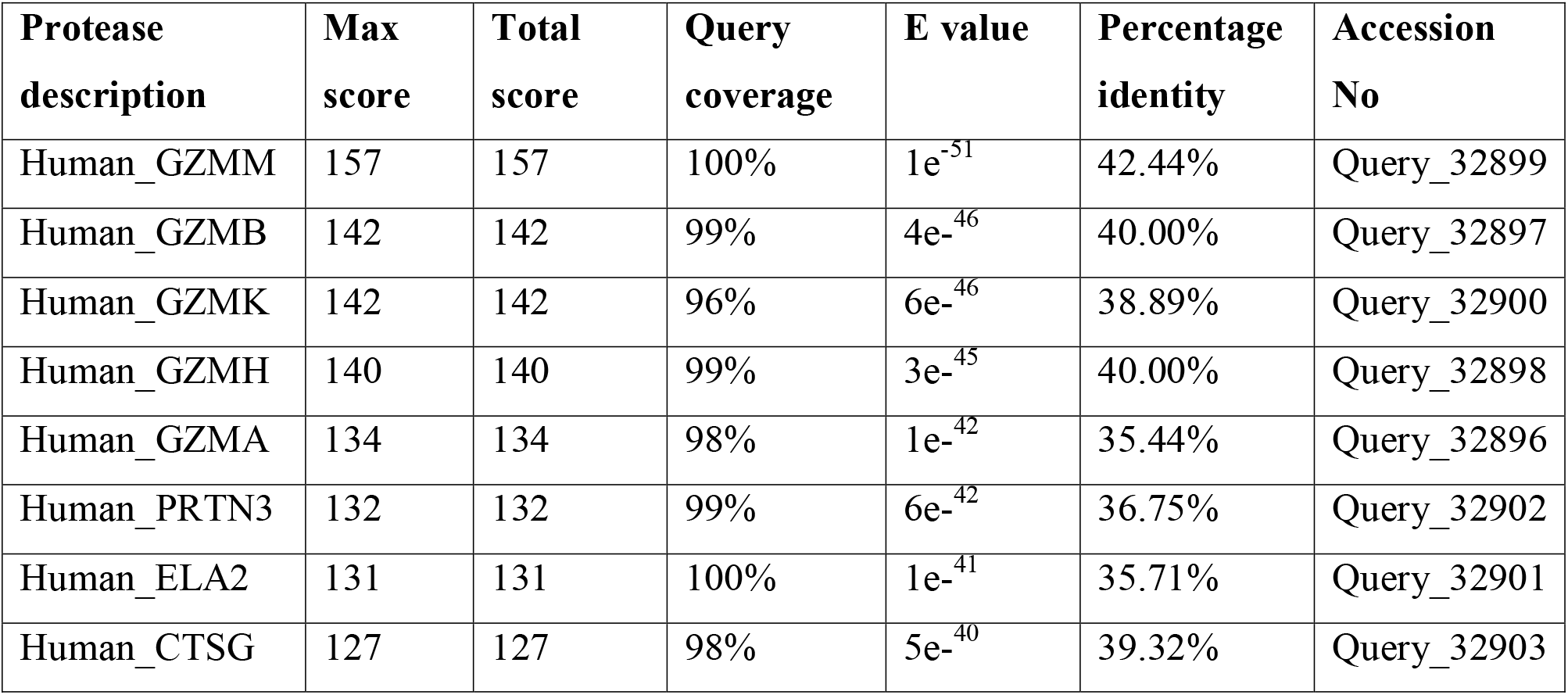
BLAST results of NSP4 against NSPs and granzymes showing percentage identity, query coverage, E value and their respective accession codes.

**Figure 1.**
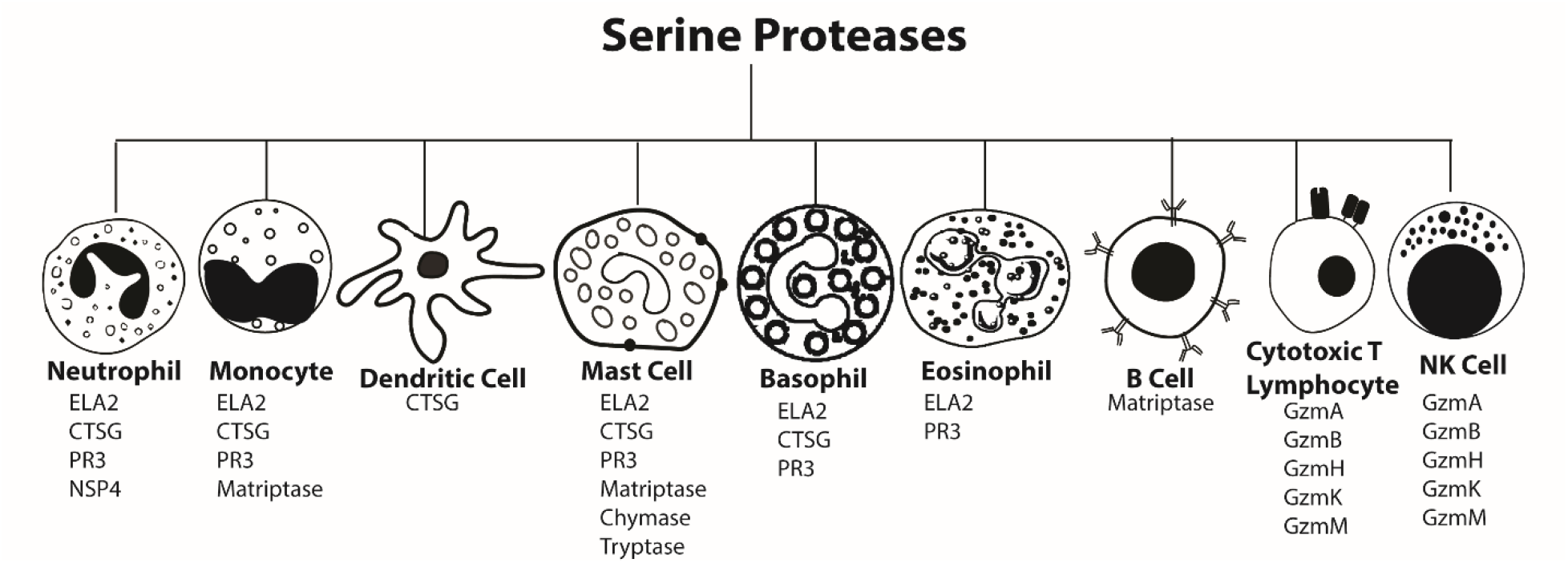
Immune cells and type of GASPIDs expressed by them.

**Figure 1.**
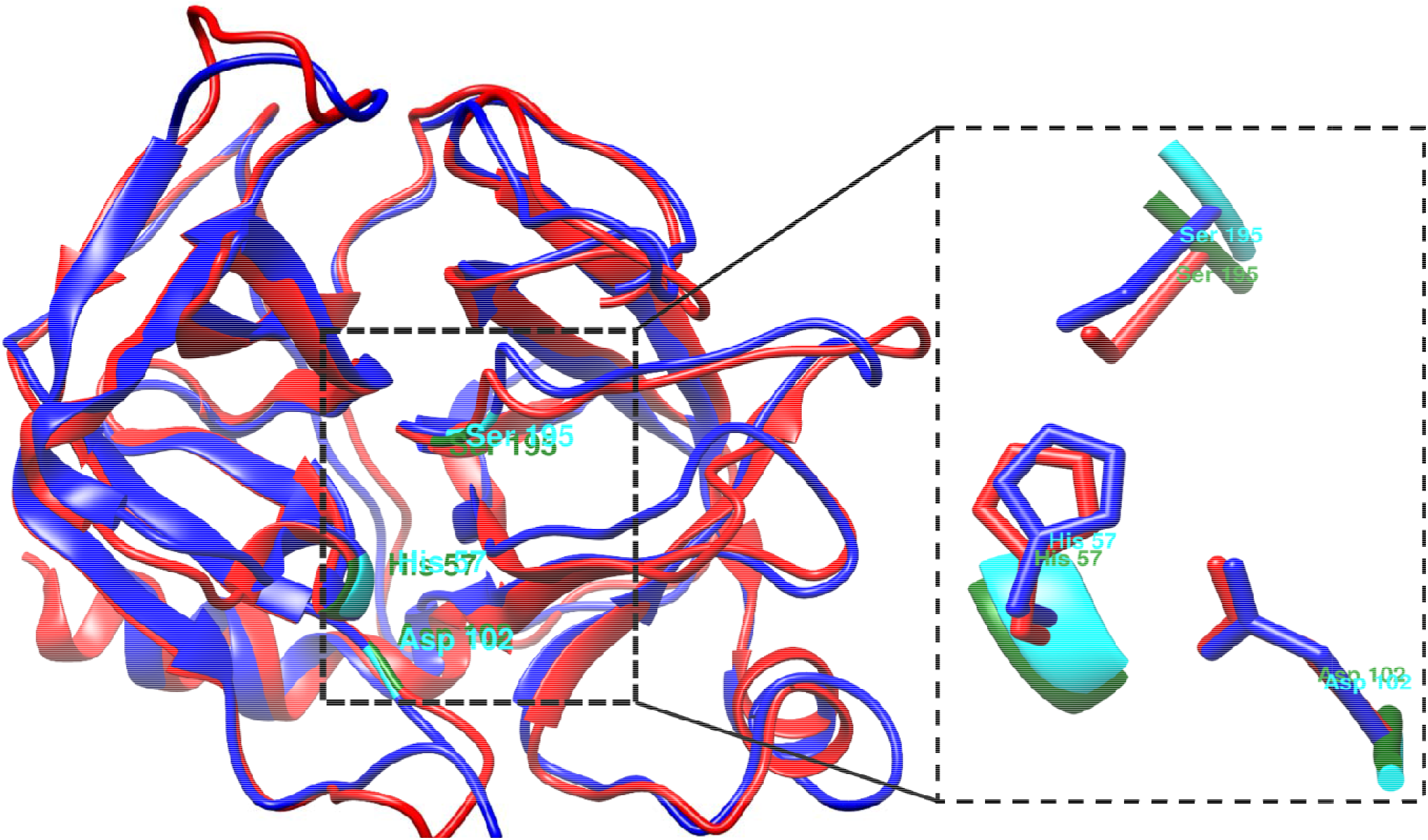
Structural representation of human NSP4 (blue) and human GZMM (red), displaying the conserved catalytic triad residues His57, Asp102, and Ser195 in a bird eye view. The active site residues of NSP4 are represented in cyan and GZMM are represented in forest green.

**Figure 2.**
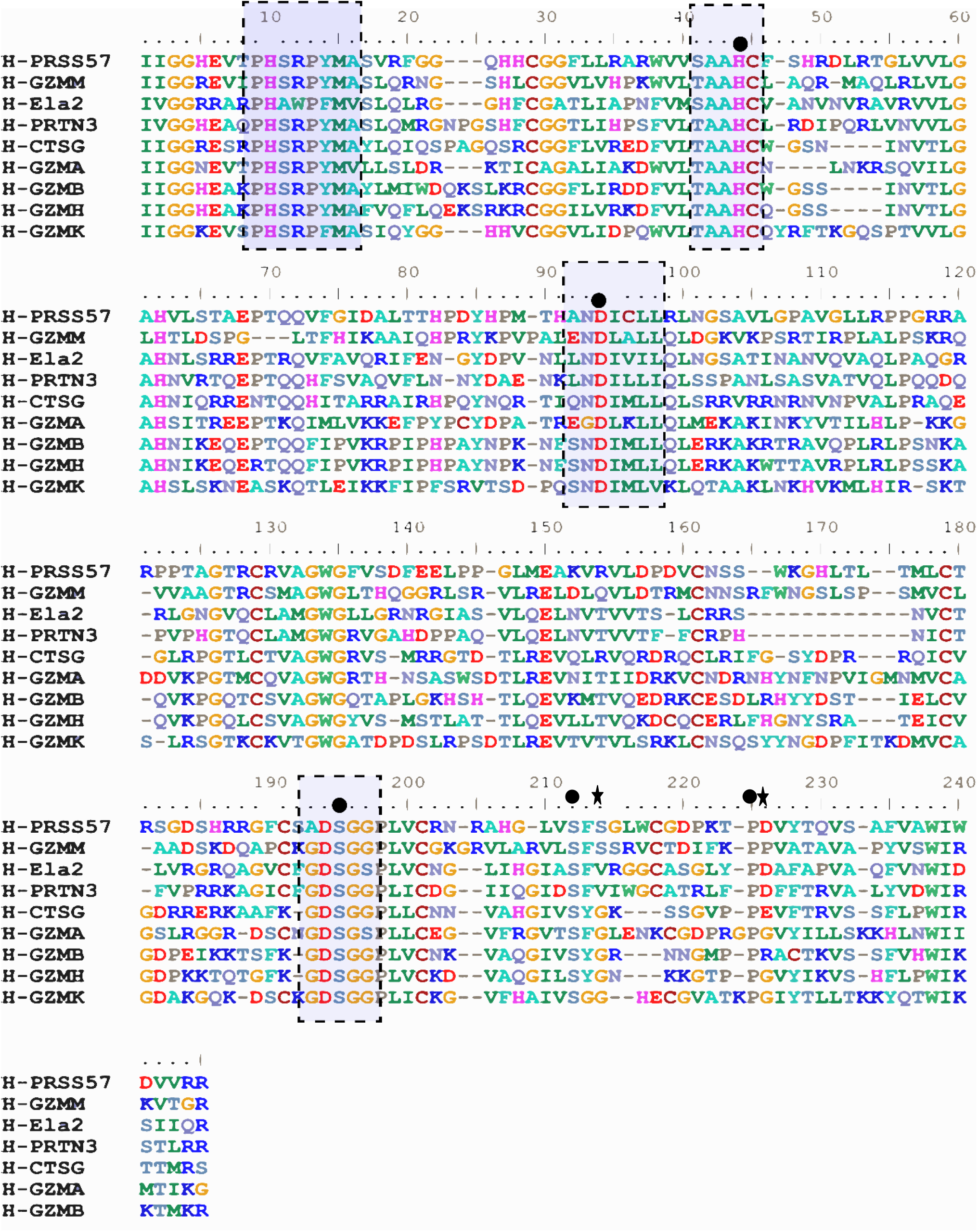
Alignment of selected GZMM and PRSS57 sequences. Highlighted regions in the square are the hallmark residues near the active site, whereas dot (•) represents highly conserved residues and at positions 214 and 225. The star (*) sign displays variation at positions 216 and 226.

BLAST was run for NSP4 against different granzymes and NSPs. Results showed a maximum identity percentage of 42.44 % with GZMM, with complete coverage, implying GZMM is closely related to NSP4 as shown in the table.

### Sequence Logo

The generated sequence logo revealed the conserved patterns and amino acid conservation in both the proteases at several positions including hallmark residues of serine proteases, as shown in Figure 3.

**Table 2.** Comparison of substrate specificity conferring residues of NSP4 and GZMM at specific positions in different species. ---represents the absence of GZMM in the frog.

| Species | Amino acid Position |  |  |  |  |  |  |  |  |  |
| --- | --- | --- | --- | --- | --- | --- | --- | --- | --- | --- |
|  | 189 |  | 190 |  | 192 |  | 216 |  | 226 |  |
|  | NSP4 | GZMM | NSP4 | GZMM | NSP4 | GZMM | NSP4 | GZMM | NSP4 | GZMM |
| **Human** | Gly | Ala | Phe | Pro | Ser | Lys | Ser | Ser | Asp | Pro |
| **Mouse** | Gly | Ala | Phe | Pro | Ser | Lys | Ser | Ser | Asp | Pro |
| **Opossum** | Gly | Ala | Phe | Pro | Thr | Lys | Ser | Ser | Asp | Pro |
| **B. Finch** | Gly | Ala | Val | Pro | Ala | Lys | Ser | Ser | Asp | Pro |
| **Croc** | Gly | Ala | Val | Pro | Ser | Lys | Ser | Ser | Asp | Pro |
| **Frog** | Gly | --- | Phe | --- | Asp | --- | Ser | --- | Asp | --- |

**Figure 3.**
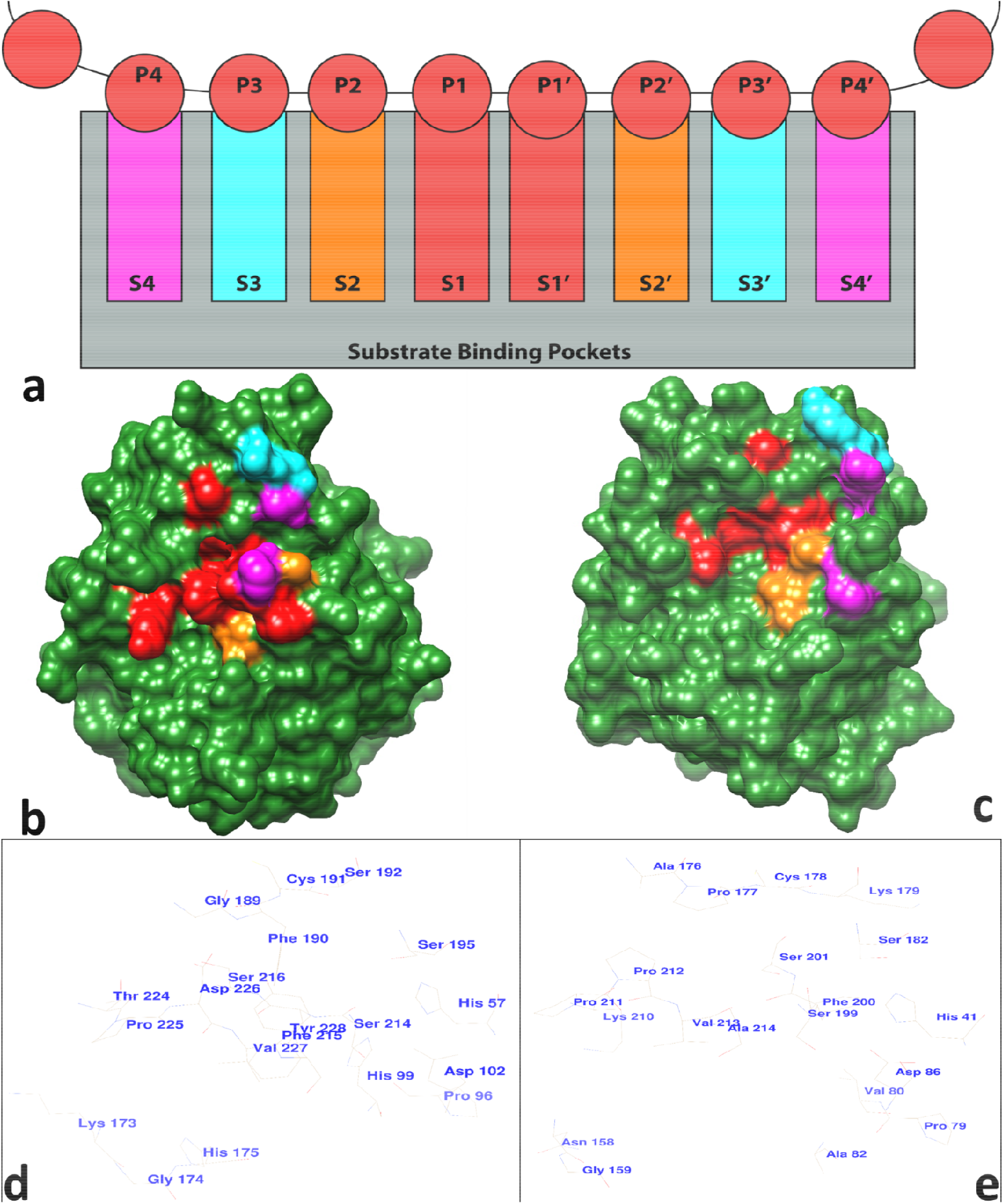
**a)** Substrate specific binding pockets of NSP4 and GZMM. **b)** Structure representation of NSP4 with substrate-specificity pockets in solid. **c)** Structure representation of GZMM with substrate-specificity pockets in solid. The substrate-specificity pockets sites are colored according to specific colors. The S1 and S1’ pocket in red; S2 and S2’ in orange, S3 and S3’ in Cyan, and S4 and S’ in Hot pink. **d)** Active site and substrate binding pocket residues of NSP4. **e)** Active site and substrate binding pocket residues of GZMM.

**Figure 3.**
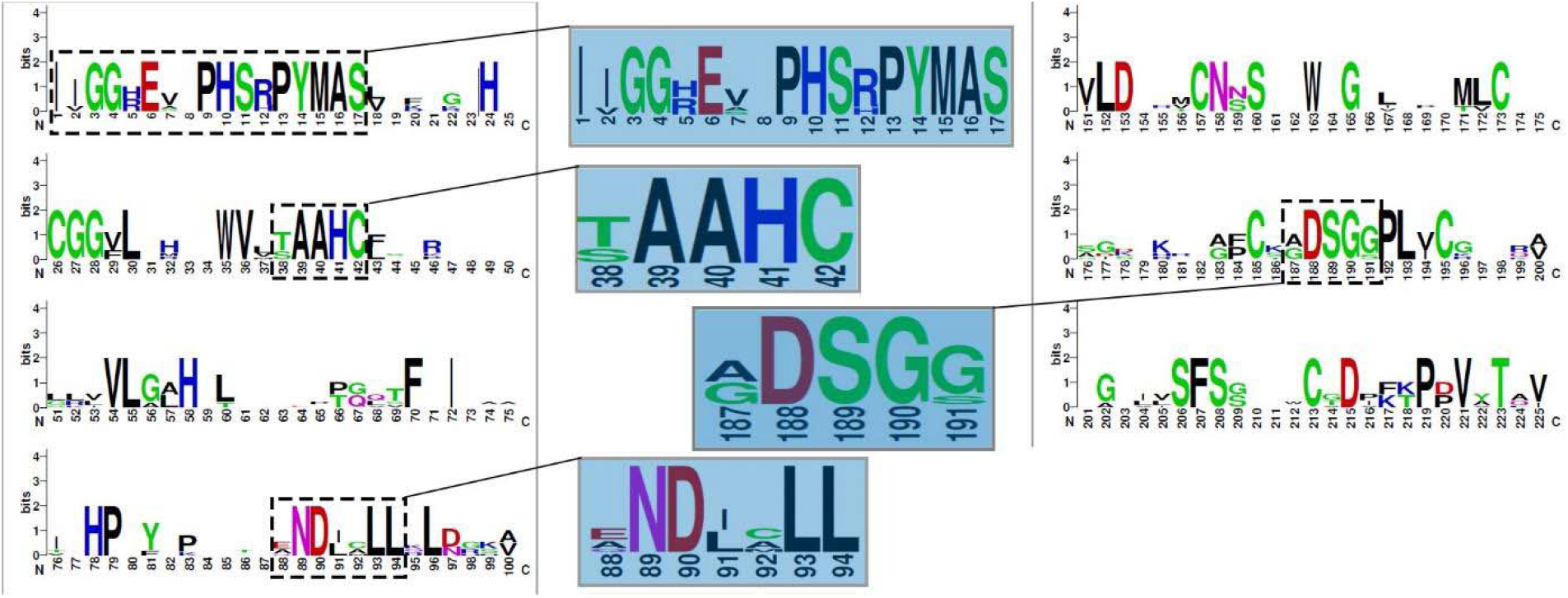
Sequence logo representation of NSP4 and GZMM showing highly conserved residues among them. The signature sequence of GASPIDs PHSRPYMAS is highly conserved among them.

### Locus map generation

The locus maps for human *Prss57* and *Gzmm* and PRSS57 were constructed. The human chromosome 19 locus consists of genes encoding 5 different GASPIDS *(Gzmm, Prss57, Prtn3, Ela2 and Cfd1, shown in red in Figure 4)* and Azurocidin *(Azu1, shown in blue in Figure 4)*, a pseudogene, accompanying the non-protease genes *(shown in black in Figure 4)*. Flanking genes of *GZMM* were found to be *Cdc34* and *Bsg*. In between *Gzmm* and *Prss57*, six non-protease genes can be seen *(Bsg, Hcn2, Polrmt, Fgf22, Rnf126 and Fstl3)*. These genes were found in the genome of placental mammals. Some non-protease genes *Palm, Misp, and Ptbp1 and Lppr3* are downstream of *Prss57* in non-placental mammals. These genes flank three NSPs (*Azu1, Prtn3 and Ela2*) and *Cfd1*, subsequently flanked by the non-protease genes *Med16 and R3hdm4*. Similar flanking genes were also identified in the mouse genome on chromosome 10 except the *Azu1* gene. In opossum on chromosome 3 *GZMM* is flanked by *Cdc34* and *Bsg*. Similar genes were observed between *Gzmm* and Prss57, as seen in humans and mice. But the later mentioned three GASPIDS (*Azu1, Prtn3 and Ela2*) were not observed in the opossum genome. Genomes of birds (Bengalese finch, Turkey, Chicken, Flycatcher) and reptile (Crocodile, turtle, snake, lizard) were searched for PRSS57 and GZMM. Both of these genes *Gzmm* (primary_assembly MUZQ01000559.1: 207,135-209,077) and *Prss57* (primary_assembly MUZQ01000339.1: 444,901-446,976) were found on separate contigs in the genome of B. Finch. In Flycatcher, no *Gzmm* was found while *prss57* was present (primary_assembly 28: 3,226,578-3,229,335). *Prss57* like gene was found in Chicken on chromosome 28 (28: 2,847,048-2,850,872) on a similar position where *Prss57* should be present but no *Gzmm* was found. Turkey genome had of both *Prss57* (primary_assembly MDVP01000004.1: 173,452-178,759) and *Gzmm* (primary_assembly MDVP01000004.1: 1,217,525-1,220,997). Similarly investigating the sequence of reptiles, the crocodile genome consisted of both *Prss57* (primary_assembly MDVP01000004.1: 173,452-178,759) and *Gzmm* (primary_assembly MDVP01000004.1: 1,239,101-1,242,577) whereas the turtle genome also had both these proteases *Prss57* (primary_assembly PRFB02000195.1: 5,600,687-5,605,700) and *Gzmm* (primary_assembly PRFB02000195.1: 3,937,156-3,942,427) on the same primary assembly but on separate contigs. Additionally, no GZMM was found in snake and lizard genomes, however, *Prss57* was present in both species.

**Figure 4.**
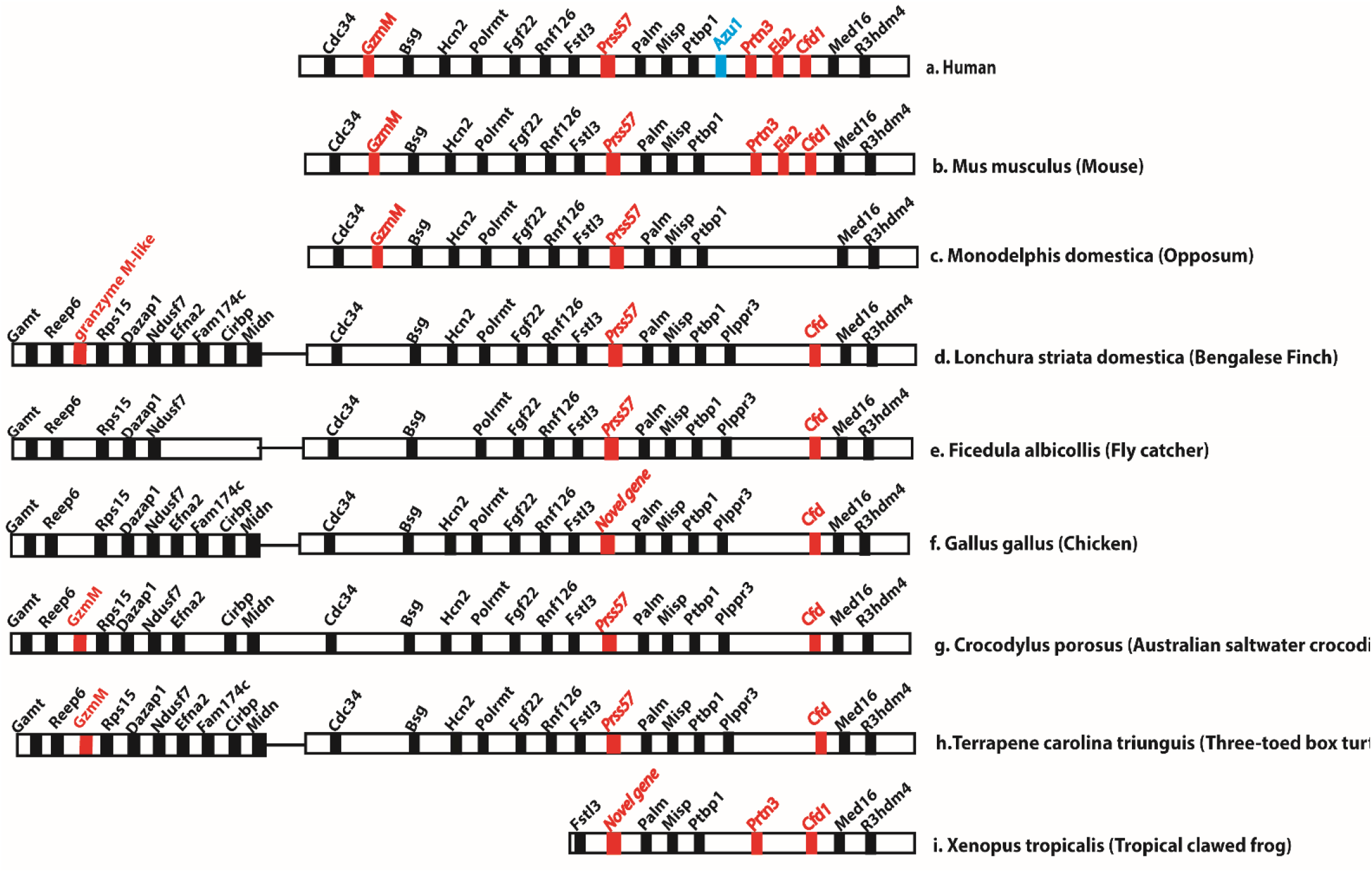
Met-ase loci in vertebrates displaying *Prss57* and *Gzmm* in a) human (chromosome 19p13), b) mouse (chromosome 10), and c) opossum (chromosome 3). d) Primary assembly of B. finch for *Prss57* (MUZQ01000339.1: 444,901-446,976) and *Gzmm* (MUZQ01000559.1: 207,135-209,077); contigs separated by a connecting bridge. e) *Prss57* primary assembly of Flycatcher (primary_assembly 28: 3,226,578-3,229,335). f) Novel *Prss57*-like gene in chicken on chromosome 28. g) Primary assembly of Australian salt water crocodile *Prss57* (MDVP01000004.1: 173,452-178,759) and *Gzmm* (MDVP01000004.1: 1,217,525-1,220,997). h) Primary assembly of three toed turtle *Prss57* (PRFB02000195.1: 5,600,783-5,605,604) and *Gzmm* (PRFB02000195.1: 3,937,257-3,942,326). i) Primary assembly of frog novel *Prss57*-like gene (Primary_assembly 1: 82,044,893-82,056,586). GASPID genes are represented in red; non-GASPID genes in black and *Azu1* in light blue. The joint in the boxes indicate the end of a contig/different assembly.

### Phylogenetic Analysis

In the first part codons for the proteases were used for constructing a consensus phylogenetic tree to look for maximum relatedness and closeness of the selected proteases. Neighbor-joining tree was constructed in MEGA. NSP4 and GZMM were found in the same clade indicating that NSP4 and GZMM are closely related proteases. Other proteases like GZMB and GZMH grouped and GZMA and GZMK were grouped. ELA2 and PR3 formed a separate clade implying relatedness. While chymotrypsin-like serine protease (CTLR) was chosen as an outgroup member. To construct tree for proteins sequences, distance matrix file was used and neighbor-joining algorithm was employed that formed a tree with a fitness score of 95.544 involving 50 taxa. The results were in accordance with the results obtain from codons. As both NSP4 and GZMM were in the same clade. Another neighbor-joining tree that involved only the C-terminal tail sequences (supplementary Figure 1) was generated only for the human GASPIDs and the results were also in accordance strongly supporting the relatedness of NSP4 with GZMM.

### Co-expression analysis

Gene names *Prss57* and *Gzmm* were given to GeneMania, a web-interface for pathway generation. The network showed co-expressing proteins as well as those that share common domains. *Gzmm*, along with other genes was found to be co-expressed with *Prss57* figure 6 (a). However, a similar generated network for *Gzmm* had no impact on *Prss57* expression figure 6 (b).

**Figure 5.**
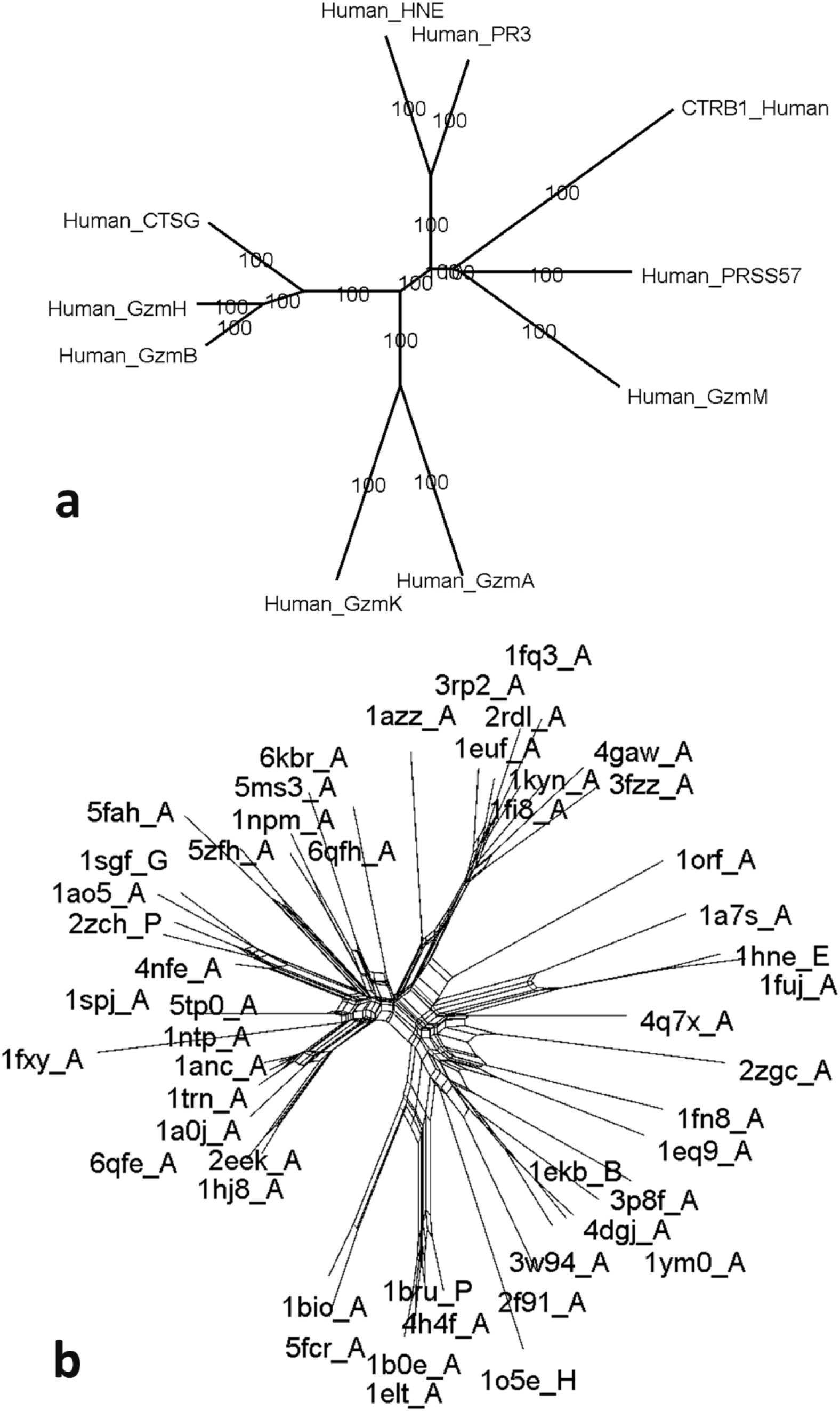
**a)** Phylogenetic tree of GASPIDs codon sequences from humans. The bootstrap replication score was set to 1000 and the confidence score was 100. **b)** Neighbor-joining tree of serine proteases by distance matrix. The proteases are named by their respective PDB IDs. PDB IDs for NSP4 and GZMM are 4q7x and 2zgc, respectively. Whereas for other GASPIDs, HNE (1hne), PR3 (1fuj), CTSG (1kyn), GZMA (1orf), GZMB (1fq3), and GZMH (4gaw).

**Figure 6:**
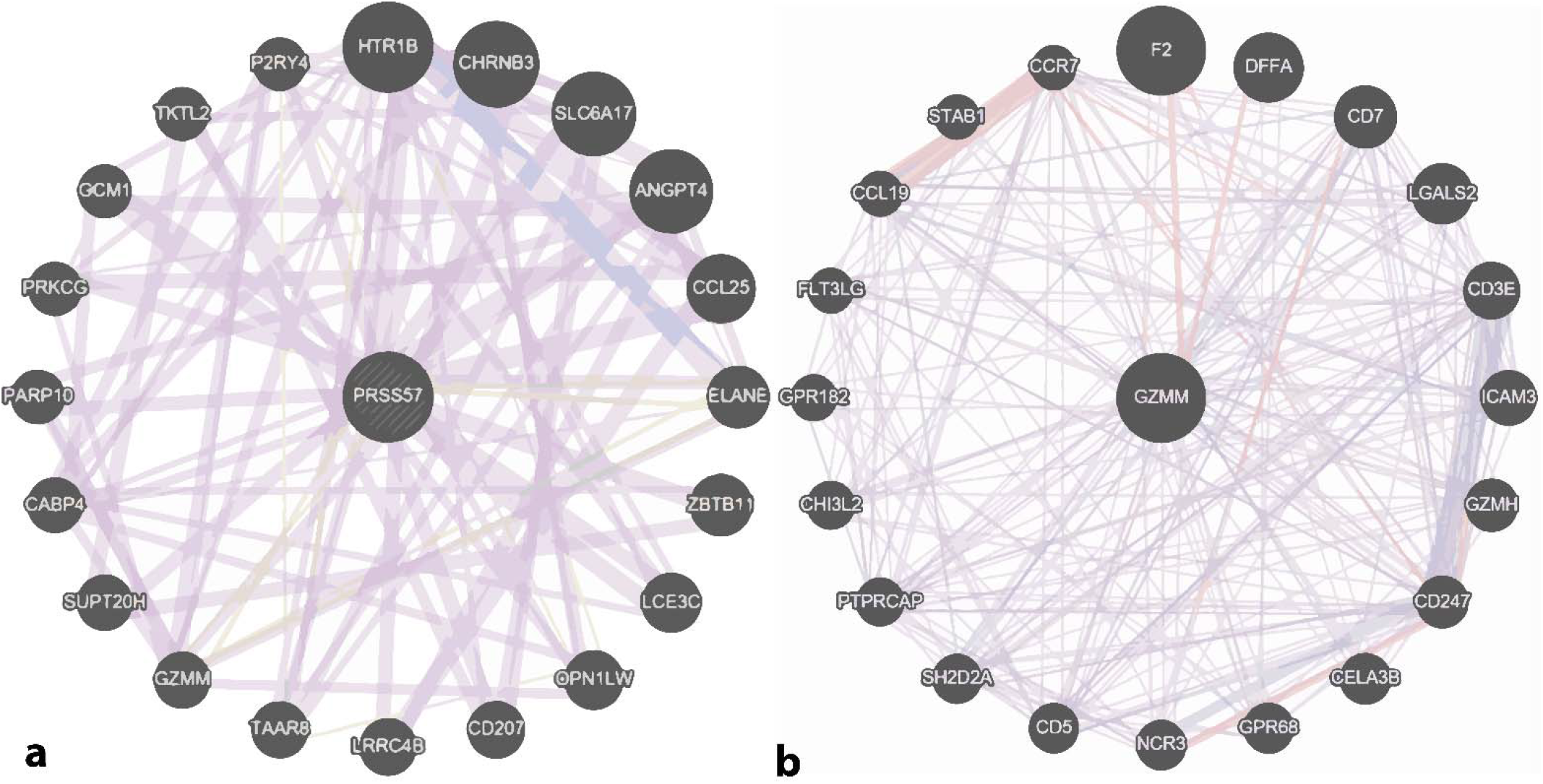
Network interaction and Co-expression analysis of **(a)** *Prss57* interacting with different proteins and **(b)** *Gzmm* interaction with different proteins. The color scheme depicts the type of interaction observed among the genes. Pink; physical interactions, purple; co-expression, orange; predicted, blue; Co-localization, cyan; pathway, green; genetic interactions, and alabaster; shared domain.

## Discussion

NSP4 being an ancestral protease [22] and sharing similar characteristics with the other NSPs remained uncovered for a long time [21]. Several studies so far have been done regarding the investigation of the substrate specificity [41,42] catalytic mechanism [15,21], and structure [22]. Recently, a study regarding the co-localization of NSPs using fluorescent probes has also been done [43]. Akula et.al described the evolutionary aspects of proteases and our results are in correspondence as NSP4 and GZMM existed in the same clade [28]. Likewise, a lot of similar investigations regarding GZMM have been studied previously [44–46]. Both these proteases are exclusively involved in conferring immunity to the individuals and reportedly possess certain similar immunological characteristics of extracellular matrix cleavage and inflammation [26,47]. We found an evolutionary relationship between *Prss57* and *Gzmm* keeping in mind that both the proteases coded by these genes are homologous and both possess more similarity as compared to the other NSPs and granzymes. All NSPs and granzymes along with active site residues have a conserved Ser at position 214 that is necessary to promote productive formation of the enzyme-substrate complex in serine proteases [48]. Additional residues for determining primary specificity at positions 216 and 226 were also observed [49]. Both NSP4 and GZMM have a conserved Ser at 216 positions but variation was observed at 226 position i.e. Asp in NSP4 and Pro in GZMM (Figure 1). However, the substrate specificity conferring residues in both the proteases are different that is obvious due to their different substrate specificity. Also, the observed codon for the conserved serine residues was TCN. NSP4 has been reported to share approximately 39 to 40% sequence similarity with its other members NSPs [21] but we found a sequence similarity of 42% between NSP4 and GZMM. NSP4 also possesses more sequence similarity with GASPIDs of CTLs and NK lineage than NSPs. However, more detailed studies are required to better understand this evolutionary tale. Similarly, GZMM presence in Met-ase locus [50] and its greater similarity with the NSP4 support the hypothesis of *Gzmm* evolution from *Prss57* by a gene duplication event somewhere back in the time that might result in a paralogue. Moreover, the expression analysis of both these genes revealed unusual findings that the *Prss57* induces *Gzmm* expression however no such patterns were observed for *Gzmm* to induce *Prss57* suggesting that as NSP4 is expressed in a very low amount in neutrophils [21,43] GZMM might be creating an immune balance and playing a role in the activation of the immune response. Nevertheless, experimental verification in the wet lab is required that will enable us to understand their correlation in a much better way. A similar co-expression analogy reportedly exists between GZMB and GZMH in the case of adenovirus infection [51,52]. Expression of GZMH sets GZMB free so that it can initiate apoptosis in infected cells [51,52]. Considering this analogy, NSP4 and GZMM might also possess a similar role in immune-related responses. We also thoroughly investigated the species for *Prss57* and *Gzmm*. Many species were found to have *Prss57* but *Gzmm* was not found although the flanking genes of *Gzmm* were present in other mammalian species. It suggest that as more sequencing is done and annotating software gets more precise many of these species might come up with GZMM and several other new proteases.

## Acknowledgement

A special thanks to Dr. Ashar Malik for useful his useful suggestions.

---

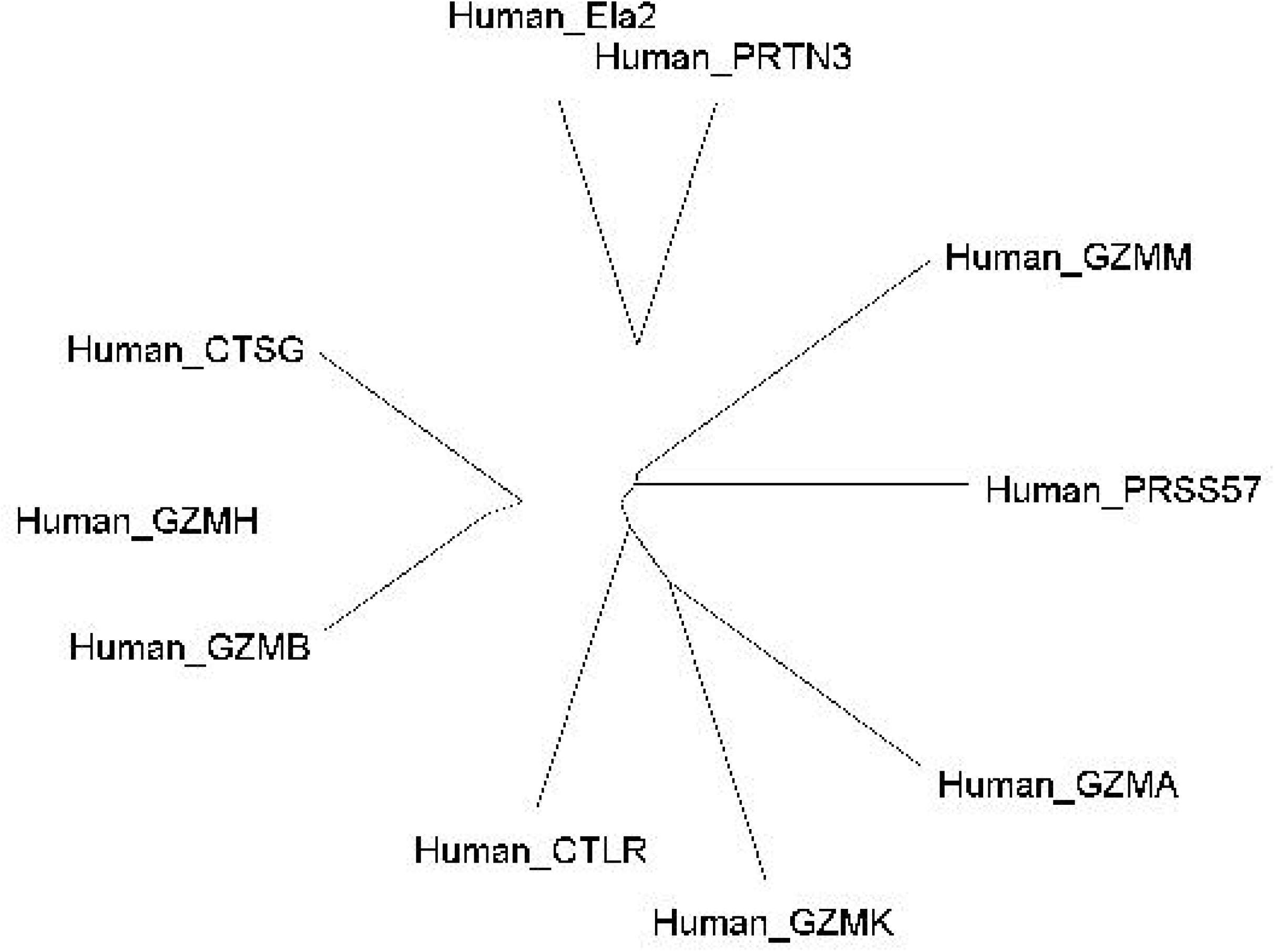

